# Antibody-Independent Antitumor Effects of Cd32A-Chimeric Receptor T Cells: Implications for Breast Cancer Prognosis and Treatment

**DOI:** 10.1101/2021.12.22.473841

**Authors:** Giuseppe Sconocchia, Giulia Lanzilli, Valeriana Cesarini, Domenico Alessandro Silvestris, Roberto Arriga, Katayoun Rezvani, Sara Caratelli, Ken Chen, Jinzhuang Dou, Carlo Cenciarelli, Gabriele Toietta, Silvia Baldari, Tommaso Sconocchia, Francesca De Paolis, Anna Aureli, Giandomenica Iezzi, Maria Ilaria del Principe, Adriano Venditti, Alessio Ottaviani, Giulio Cesare Spagnoli

**Author notes:** **Corresponding Author:** Giuseppe Sconocchia, Institute of Translational Pharmacology, National Research Council of Italy, Via Fosso del Cavaliere 100, 00133, Rome, Italy. These authors have equally contributed to this work (Alessio Ottaviani, Giulio Cesare Spagnoli). Actual address: Takis Biotech, via di Castel Romano 100, Rome Italy (Roberto Arriga).

## Abstract

Fcγ RIIA (CD32A) and their ligands, including the immunoglobulin Fc fragment and pentraxins, are key players in a variety of innate immune responses. Still unclear is whether additional ligands of CD32A do exist. The objective of this study is to demonstrate that CD32A-chimeric receptor (CR) can be utilized for the identification of CD32A cell surface ligand(s). Among fifteen cancer cell lines tested, CD32A-CR T cells recognized three of breast cancer (BC) including the MDA-MB-468 and one colorectal carcinoma (HT29) in the absence of targeting antibodies. Conjugation of sensitive BC cells with CD32A-CR T cells induced CD32A polarization and down-regulation, CD107 release, and mutual cell elimination *in vitro*. Conversely, normal fibroblasts and myoblasts were not affected while normal HUVEC cells promoted CD32A down-regulation. CD32A-CR T cell activity was not inhibited by human IgGs or human serum, but; it was rather enhanced by cetuximab antibody. RNAseq analysis of sensitive vs resistant BC cells identified a fingerprint of 42 genes predicting the sensitivity of BC cells to CD32A-CR T cells and their association with favorable prognostic significance in advanced BC patients. Our data also identify ICAM 1 as a major regulator of CD32A-CR T cell-mediated cytotoxicity. Finally, CD32A-CR T cell administration protected immunodeficient mice from subcutaneous growth of MDA-MB-468 cells in the absence of tumor-specific antibodies. These data indicate that CD32A-CR can be utilized for the identification of (1) cell surface CD32A ligand(s); (2) rational therapeutic strategies to target BC; and (3) novel transcriptomic signatures prognostically relevant for advanced BC patients.

## INTRODUCTION

Innate immune cells represent the first line of defense against pathogens, virally infected and cancer cells. Innate immune cells comprise a variety of myeloid and lymphoid cell (ILC) types. Myeloid cells include granulocytes, monocyte/macrophages, and dendritic cells (DC)^1,2^. These cells regulate the host’s inflammatory responses by releasing cytokines^3,4^, performing phagocytosis ^1^, mediating direct cytotoxic functions ^5^, and tissue repair following ischemic and thermal injuries ^6–8^. Lymphoid cells include natural killer (NK) cells and ILC mainly involved in tissue homeostasis and mucosal immunity. Among innate immune cell receptors, Fc gamma receptors (FcγRs) play key roles in the regulation of NK, ILC ^9^, DC, and myelomonocytic cell functions ^9^. By mediating antibody-dependent cellular cytotoxicity (ADCC) phagocytosis (ADP), and cytokine release, they are critically involved in the response to infectious challenges and the pathogenesis of inflammatory, infectious diseases, as well as in tissue damage and remodeling ^4,5,10–13^.

FcγRs include three distinct sets of molecules: the activating FcγRI (CD64), FcγRII (CD32A, CD32C), and FcγRIII (CD16A and CD16B), and the single inhibitory FCγRII (CD32B). FcγRs bind γ immunoglobulins (Igs) with different affinities ^12,14–16^. While CD64 is a high-affinity receptor expressed on IFNγ-activated granulocytes^4^, monocyte/macrophages, and DCs ^17^, CD32 and CD16 are polymorphic with low binding affinity for monomeric IgG.

Fcγ activating receptors transduce signals through the Immunoreceptor tyrosine-based activation motif (ITAM). CD64 and CD16A consist of dimers composed of a ligand-binding chain and a signal subunit γ chain containing ITAM. Instead, CD32A and CD32C are monomeric directly transducing activation signals through the ITAM located in the intracellular tail ^12^.

CD16A is the most important FcγR mediating ADCC. It is expressed on NK cells and small subsets of monocytes and NK T cells ^9,18,19^. NK cells efficiently mediate ADCC against tumor cells. However, despite the high expression of NK cell ligands including MICA/B, in the microenvironment of solid cancers, NK cell infiltration is rarely detectable and has no direct prognostic impact ^20–24^. In contrast, CD8+ T cells are promptly identified in the solid tumor microenvironment. These cells are associated with a favorable clinical course of the disease and are easy to expand *in vitro*.

Following the demonstration that rituximab could redirect CD16A engineered T cells to kill B lymphoblastic cells ^25^, we and others have independently produced a variety of CD16A chimeric receptors (CR) T cells to target solid tumor cells ^26–28^. Also, NK cells have demonstrated weak ADCC, when utilized in combination with IgG2 mAb including panitumumab, while IgG2 mAb elicits ADCC in CD32A positive myeloid cells ^29^. Therefore, to overcome CD16A limitation, CD32A-CR has also been produced ^28 30–32^. They share, with classical CAR, the intracellular tail, resulting from the fusion of the transmembrane CD8α with the co-stimulatory molecule CD28 linked to the T cell CD3ζ chain (CD32A/CD8a/CD28/ζ). In contrast, the extracellular CAR single-chain variable fragment (scFv), which recognizes the target tumor-associated antigen (TAA), has been replaced with the extracellular portion of CD32A. CD32A-CR T cells bind IgG2 (panitumumab) even in the presence of human serum while CD16A-CR does not. In addition, panitumumab successfully redirects CD32A-CR T cells toward cancer cells overexpressing EGFR ^28,31,32^.

Other than Igs, FcγR binds pentraxins, a family of soluble proteins involved in opsonization, phagocytosis, and complement activation. The best-known pentraxins are the C-reactive protein (CRP) and serum amyloid component (SAP)^33,34^. Most interestingly, however, previous evidence suggests that at least CD16A mediates NK cell cytotoxicity in the absence of cell surface antigen-bound antibodies consistent with its ability to directly recognize poorly defined, cell surface ligand(s) expressed by tumor target cells ^35^.

To date, the existence of CD32A cell surface putative ligand(s) is still unknown. In this study, utilizing CD32A-CR as a biosensor, we provide evidence of the existence of CD32A surface ligand(s) on cancer cells and that CD32A-CR T cells can be used as a therapeutic and prognostic tool for early clinical studies. The presented work is likely to enhance the basic and translational field of CD32A investigation behind ADCC and ADP.

## RESULTS

### CD32A-CR promotes specific T cell surface recognition of BC cells

To provide evidence of the existence of putative CD32A cell surface ligand(s) on cancer cells, we tested the ability of tumor cells from 15 established cell lines to trigger CD32A-CR T cells (table1). Incubation with triple-negative breast cancer (TNBC) cells, MDA-MB-468 and MDA-MB-231, induced significant CD32A-CR down-regulation in 78.7±7% and 81.3±8% of engineered T cells respectively (fig.1A). Concomitantly, CD107A mobilization was also observed. Representative contours and cumulative data are reported in figure 1A. Conjugation of MDA-MB-468 and MDA-MB-231 cells with CD32A-CR T cells induced CD32A-CR T cell down-regulation. To provide improved visualization of the effects elicited by the interaction between BC and engineered T cells, we performed a confocal microscopy analysis. CD32A-CR T cells (fig.1B, panel a) and MDA-MB-468 (fig1B, panel b) were indirectly stained with anti-CD32 mAb (green) and anti-EGFR mAb (red) respectively. Incubation of CD32A-CR T cells with MDA-MB-468 cells resulted in the formation of lytic synapsis including firm adhesion (fig.1B panel c), CD32A-CR polarization and down-regulation (fig.1B panels d and e), and structural cell aberrations (fig.1B panel f) including cell blebbing formation in MDA-MB-468 cells (red blebs) and following MDA-MB-468 and CD32A-CR T cell membrane fusion (yellow blebs). These results indicate that CD32A-CR senses natural ligand(s) on TNBC cells.

**Table1.**
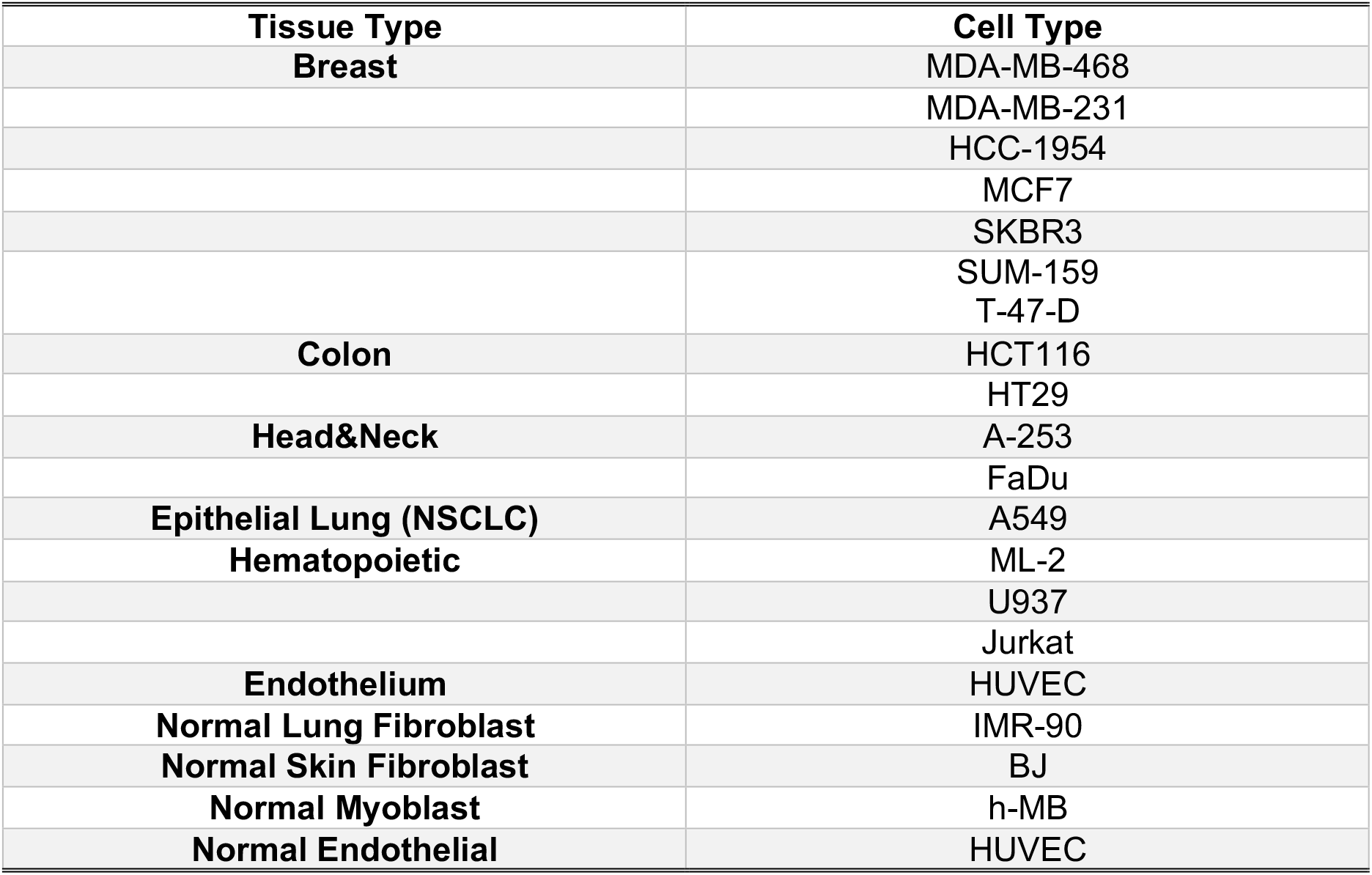
List of cells screened for induction of CD32A-CR downregulation in CD32A-T cells

| **Tissue Type** | **Cell Type** |
| --- | --- |
| **Breast** | MDA-MB-468 |
|  | MDA-MB-231 |
|  | HCC-1954 |
|  | MCF7 |
|  | SKBR3 |
|  | SUM-159 |
|  | T-47-D |
| **Colon** | HCT116 |
|  | HT29 |
| **Head&Neck** | A-253 |
|  | FaDu |
| **Epithelial Lung (NSCLC)** | A549 |
| **Hematopoietic** | ML-2 |
|  | U937 |
|  | Jurkat |
| **Endothelium** | HUVEC |
| **Normal Lung Fibroblast** | IMR-90 |
| **Normal Skin Fibroblast** | BJ |
| **Normal Myoblast** | h-MB |
| **Normal Endothelial** | HUVEC |

**Figure 1.**
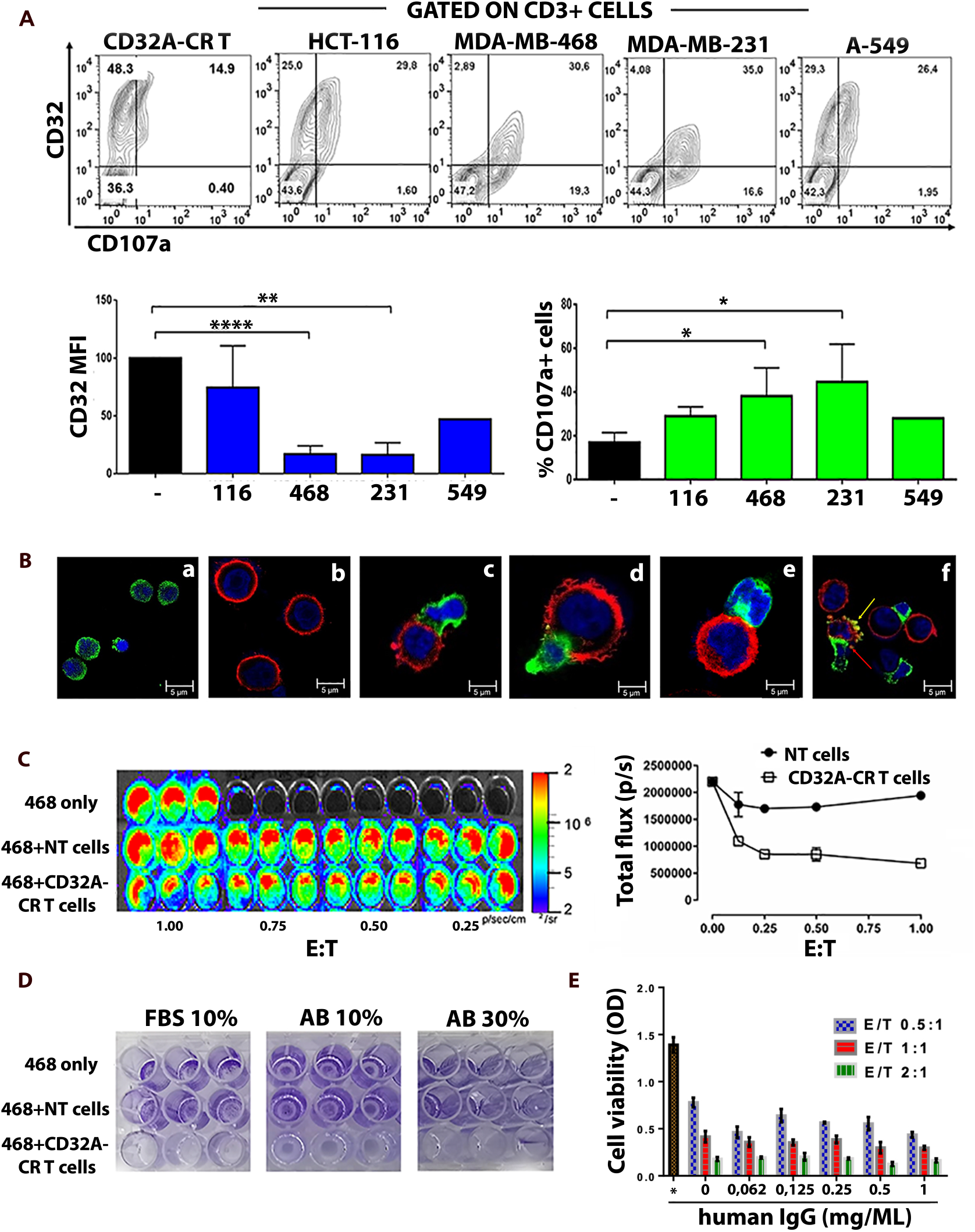
CD32A-CR T cells exert direct anti-tumor activity against TNBC cells. **A**: upper panel, CD32A-CR T cell engagement of TNBC cells (MDA-MB-468 and MDA-MB-231) resulted in CD32A-CR down-regulation and CD107a release. CD32A-CR T cells alone or following incubation with cancer cells, at an E: T ratio of 2:1, were incubated for 1hr at 37°C with a FITC-mouse anti-human CD107a. Following 1hr incubation, cell mix was incubated for 4hr at 37°C with monesin (2mM) and washed. Cells were then stained with a PE-mouse anti-human CD32 mAb and APC-mouse anti-human CD3 mAb and analyzed by flow cytometry. CD32A-CR T cells were identified by gating on CD3+cells. Lower panel: cumulative analysis of CD32Adown-regulation (left) and CD107a release (right) by CD32A-CR T cells of 3 donors as induced by cancer cells. X-axes: abbreviations of the cancer cell lines indicated in the upper panels; MFI: mean fluorescence intensity **B:** confocal imaging of CD32A–CR T [(green)a] and MDA-MB-468 cells [(red)b] following a 2hr incubation (E:T 2:1) at 37°C shows the formation of lytic synapsis consisting of firm adhesion (c), CD32A-CR dipolar localization (d), internalization (e), and tumor cell and lymphocyte blebbing (f). Red and yellow arrows, in panel f, indicate blebs generated from damaged tumor cells and both, effector and cancer cells respectively. Images were acquired using Leica confocal microscope at magnification, X 63 (oil immersion). **C:** image (left) and quantification (right) of light emission generated by luciferase-expressing cells through ATP-dependent conversion of luciferin to oxyluciferin used as a non-lytic, real-time cell viability assay. MDA-MB-468 cells were incubated for 72hr, at 37°C, with or without CD32A-CR T cells at the indicated E:T ratios. NT-T cells were used as a control. **D**: MDA-MB-468 cells were incubated with engineered T cells or NT-T cells at an E:T ratio of 2:1 as described in panel C. Then, T cells and MDA-MB-468 non-adherent (dead) cells were removed while adherent cells were stained with crystal violet. **E**: triplicates of MDA-MB-468 cells were incubated for 72hr at 37°C at different E:T ratios with or without scalar concentration of IgG as indicated. Then, non-adherent cells were removed and viable cells were assessed by the MTT assay as described in the methods. Data are representative of three experiments obtained with similar results. Values are expressed as mean ±SD. *P < 0.05; **P < 0.01; ****P < 0.0001

### CD32A-CR mediates a direct T cell anti-cancer activity

To assess whether CD32A-CR triggering might result in direct anti-tumor activity, MDA-MB-468^LUC^ (luciferase-expressing) cells were incubated in the presence or absence of CD32A-CR T cells while non-transduced (NT)-T cells were utilized as negative controls. Following 72hr incubation at 37°C, engineered T cells significantly reduced the biosignal produced by MDA-MB-468^LUC^ cells even at a very low E:T ratio (0.125:1) while the non-transduced (NT)-T cells were completely ineffective (fig.1C).

Since human IgG can bind CD32A-CR^32^, binding of CD32A-CR to its putative ligand(s) on MDA-MB-468 may be hindered in the presence of serum Igs. Therefore, MDA-MB-468 cells were incubated for 72hr, at 37°C in the absence or presence of NT-T or engineered T cells with or without human serum (fig.1 panel D) or in a complete medium supplemented with scalar concentrations of human IgGs (fig.1 panel E). Interestingly, CD32A-CR T cell anti-cancer activity was neither affected by human AB serum (10-30% concentration), as shown by the reduced crystal violet staining of adherent MDA-MB-468 cells (fig.1D), nor by human IgGs (fig.1E). Similar results were obtained when the FcR blocking buffer was utilized (data not shown). These results strongly suggest that the direct anti-tumor effect of CD32A-CR is IgG independent and that IgG and putative ligand(s) have distinct binding sites.

To demonstrate that the direct anti-TNBC activity of CD32A-CR T cells implies T cell-mediated cytotoxicity, CD32A-CR T cells were incubated for 3hr with MDA-MB-468 cells at a 1:1 E:T ratio. Engineered T cells induced direct apoptosis/necrosis of MDA-MB-468 cells (fig.2A, lower-left panel) while NT-T cells did not (fig.2A, middle-left panel). In addition, the cytotoxic activity of cetuximab (fig.2A, upper-right panel) was minimally affected by NT-T cells (fig.2A, middle-right panel) while it was significantly enhanced by CD32A-CR T cells (figure2A, lower-right panel). These data indicate that while improving therapeutic mAb effects, CD32A-CR also triggers antibody-independent T cell-mediated cytotoxicity.

**Figure 2.**
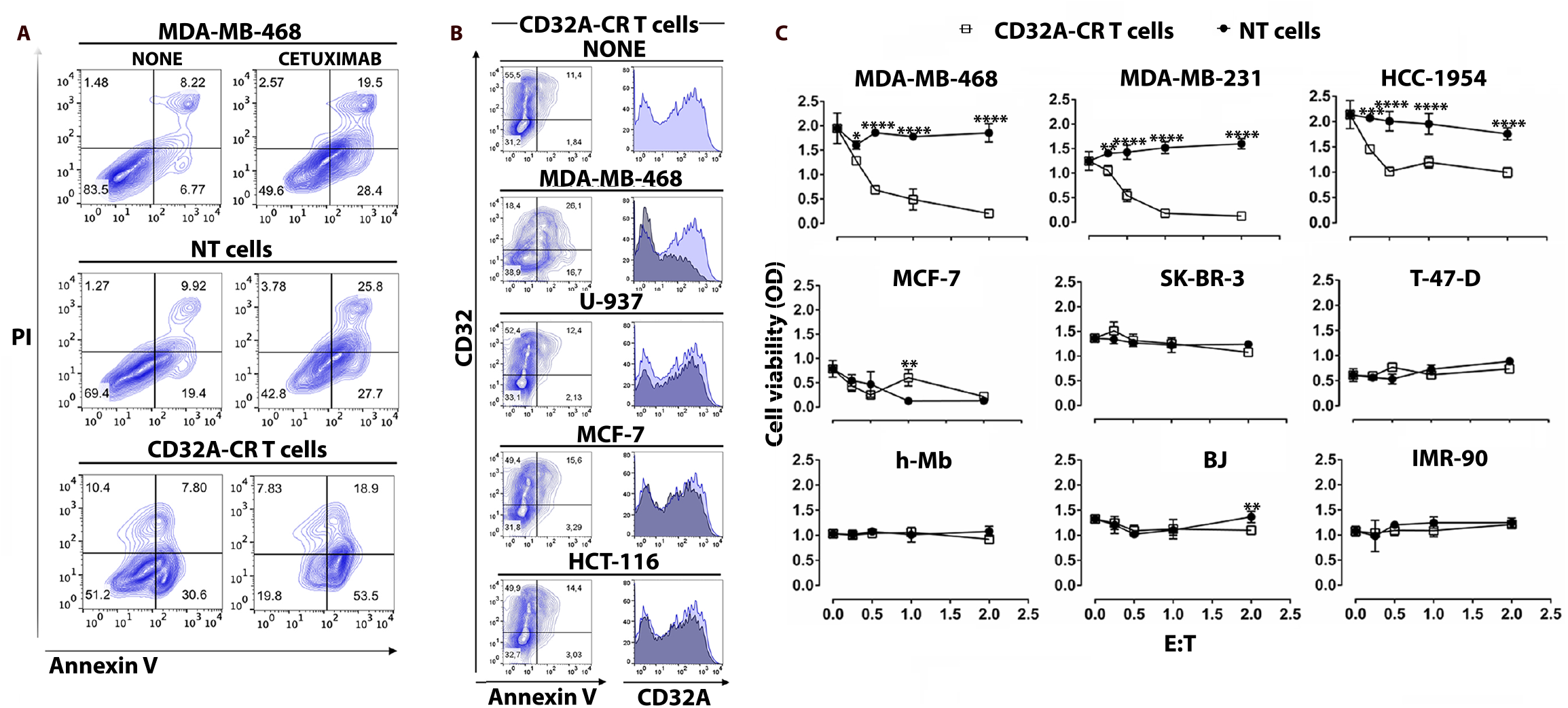
CD32A-CR T cell cytotoxicity and target cell specificity. **A and B:** following a 4hr incubation of CD32A-CR T cells or NT T cells, in the presence or absence of the indicated cell lines, were harvested, stained with APC-anti-CD3 mAb, propidium iodide (PI), and/or FITC-annexin V, and analyzed by flow cytometry. **A**: contour plots showing the cytotoxic activity of CD32A-CR T cells on MDA-MB-468 cells, in the presence or absence of cetuximab, were evaluated by gating on CD3 negative cells. **B**: apoptosis induced by the indicated cancer cells in CD32A-CR T cells was analyzed by gating on CD32A-CR positive cells. Data are representative of five experiments independently performed. **C**: preferential anti-cancer activity of CD32A-CR T cells was not limited to TNB cells. BC cells or normal cells were incubated for 12hr, at 37°C in the presence or absence of CD32A-CR T cells at the indicated E:T ratios. NT-T cells were used as a control. Cell viability was evaluated by MTT assay. Values are expressed as mean ± SD.

Recent studies have provided evidence that upon conjugation with leukemia or solid tumor cells, NK cells may undergo cellular aberrations including CD16A downregulation and apoptosis leading to mutual effector-target cell elimination ^20,21,36^. We asked whether similar aberrations can also be detected upon conjugation of CD32A-CR T cells with TNBC cells. Figure 2B shows that following a 3-hr incubation, MDA-MB-468 induced CD32A-CR T down-regulation, and CD32A-CR T cell apoptosis/necrosis, thereby, suggesting that effector and target cells undergo mutual elimination in the absence of mAbs.

### CD32A-CR T cell-mediated cytotoxicity is not limited to TNBC cells but does not involve primary myoblasts and fibroblasts

The molecular classification of human BC relies on the expression of the hormone receptors (HR) and epidermal growth factor-2 (HER2) on cancer cells. Hence, four molecular subtypes have been identified namely luminal A (HR+/HER2-), luminal B (HR+HER2+), HER2 enriched, and triple-negative (HR-/HER2-)^37^. Based on this information and the previously shown data, we investigated whether the cytotoxic activity of the CD32A-CR T cells was merely restricted to TNBC cell lines.

Therefore, engineered T cells were tested against 4 additional non-TNBC cell lines including HCC-1954 and SK-BR-3 (HER2 enriched) MCF-7 and T-47-D (luminal A) ^38^ and three primary myoblasts (h-MB), skin fibroblasts (BJ), and lung fibroblasts (IMR-90) human cell lines (fig.2C). The viability of the HCC-1954 cells was significantly affected by CD32A-CR T cells (fig.2C, upper–right panel) although not as much as that of TNBC cells (fig.2C, upper left and middle panels) while the control NT T cells were fully ineffective. It must be noted that the anti-tumor activity of CD32A-CR T cells was impressive and even detectable at an E:T ratio lower than 0.5:1. In contrast, MCF-7, SK-BR-3, and T-47-D were resistant to CD32A-CR T cells (fig.2 middle panels). Most interestingly, neither engineered T cells nor NT T cells impacted the viability of the three primary, non-cancerous, cell lines (fig.2C lower panels). These data suggest that the activity of CD32A-CR T cells is not restricted to TNBC cell lines, but; it does not affect normal myoblasts and skin and lung fibroblasts.

### RNA-sequencing analysis identified transcriptomic signatures of 73 genes associated with susceptibility of breast cancer cells to CD32A-CR T cells cytotoxicity

To gain insights into the molecular mechanisms underlying differential susceptibility of BC cells to CD32A-CR T cells and to identify relevant transcriptional signatures, publicly available RNA sequences (RNA-Seq) were interrogated. RNA-Seq from MDA-MB-468, MDA-MB-231, and HCC-1954 sensitive and MCF-7, SK-BR-3, and T-47-D resistant cells was analyzed in detail. Out of 60.469 genes tested, 1.418 were found to be differentially expressed in sensitive and resistant BC cell lines. Among these genes, 902 were upregulated and 516 were downregulated in sensitive vs resistant cells as summarized in the volcano plot (fig.3, panel A).

**Figure 3.**
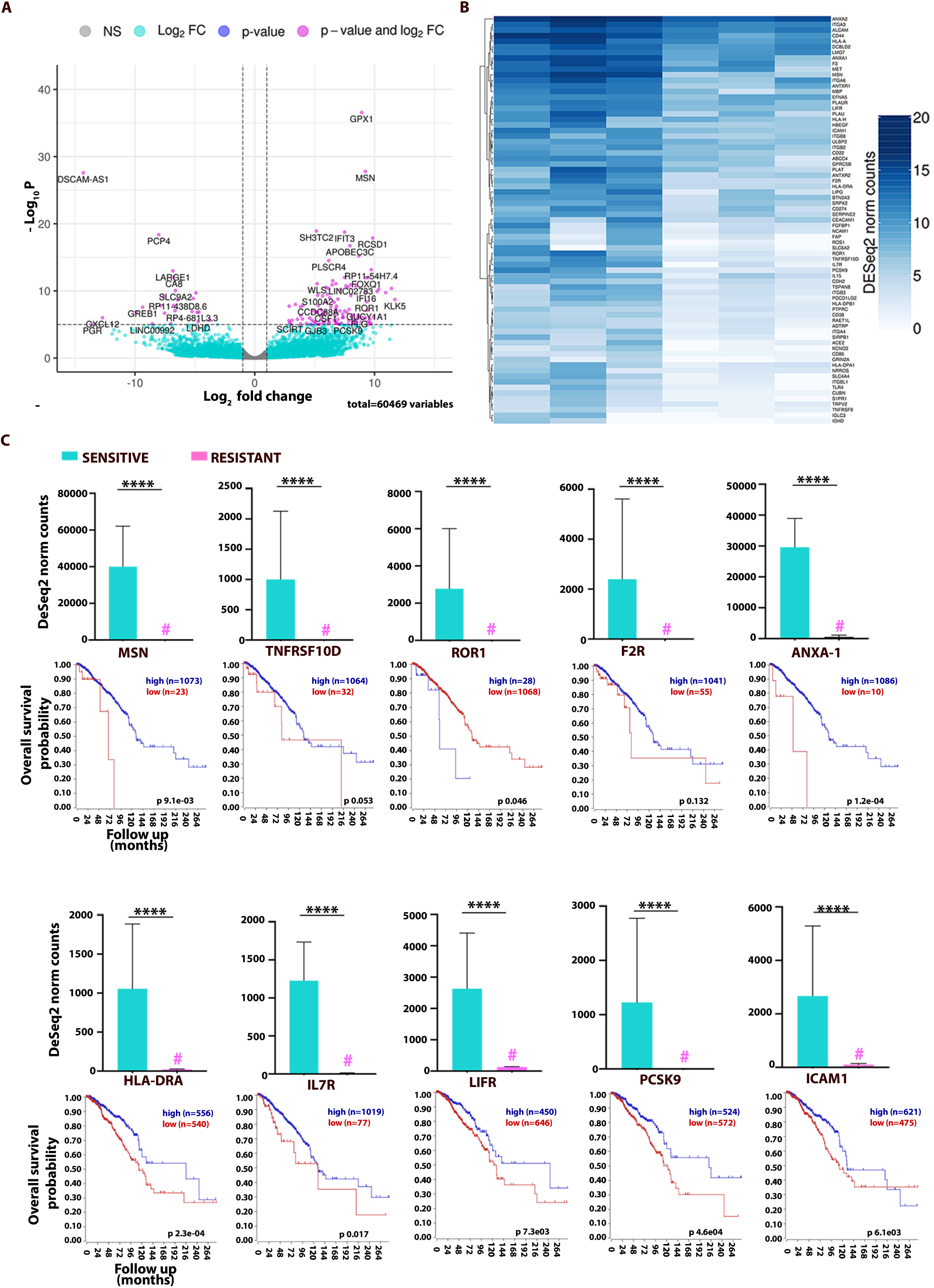
Sensitivity to CD32A-CR T cells cytotoxicity is associated with upregulated expression of genes correlating with favorable prognosis in advanced BC patients. **A**: volcano plot showing differential transcriptome of BC cells with different sensitivity to CD32A-CR T cell-mediated cytotoxicity analyzed in terms of significance levels and fold changes in gene expression, as estimated from DESeq2 analysis of RNA-Seq data. A total of 60469 genes were analyzed and 1418 were found differentially expressed in sensitive and resistant BC cell lines. Among these genes, 902 were upregulated and 516 down-regulated in sensitive vs resistant BC cells. The magenta points represent genes with log2FC >|2| and p.adj <10e-6 **B**: heatmap showing the expression levels of 73 genes encoding cell surface proteins, selected based on the Gene Ontology Cellular component category significantly upregulated in sensitive vs resistant BC cells. **C**: the 10 genes with the most significant p.adj were selected and their expression values were plotted as DESeq2-normalized counts (mean ± standard error). For each gene, a Kaplan-Meier curve, analyzing their prognostic significance as observed in 1.097 breast invasive carcinomas from TCGA, is shown (https://hgserver1.amc.nl/cgi-bin/r2/main.cgi). The expression levels of 7 out of 10 genes are significantly associated with favorable prognosis in patients with breast invasive carcinomas. **** p.adj ≤ 0.0001

To identify the key factor(s) conferring sensitivity to BC cells, we focused our attention on genes significantly upregulated in the 3 sensitive BC cell lines. To further filter involved genes, this list was annotated based on the Gene Ontology Cellular component category, thus allowing the selection of 73 genes coding for cell surface proteins (fig. 3B). Then, the clinical significance of the overexpressed genes associated with the BC cell susceptibility to CD32A-CR T cells was then assessed by using a clinically well-documented TCGA advanced BC-RNA-seq database R2: Genomics Analysis and Visualization Platform (htpp://r2.amc.nl). We first analyzed the top 10 representative genes identified in the sensitive cells with the highest differential expression compared to that of resistant BC cells. Figure 3C shows that the expression level of these genes was significantly higher than that detected in the BC resistant cells (bar graph panels). Importantly, a high expression of 7 of these 10 genes was significantly associated with improved survival in advanced BC patients. Only ROR 1 gene appeared to be associated with a poor prognosis. Subsequently, we extended this investigation to the other differentially expressed genes. Table 2 shows a direct association between a CD32A-CR T cell susceptibility and additional 35 genes bringing the number of genes associated with favorable prognosis outcomes up to 42 (table 1). Interestingly, these 42 genes were homogeneously expressed in 3 out of 3 susceptible BC cells while the remaining 31 genes were more heterogeneous (fig.3B). These data suggest that the sensitivity of BC cells to CD32A-CR T cells may predict a favorable prognosis in advanced BC prompting a previously unsuspected anti-tumor role in BC immunobiology. On the other hand, the prognostic significance of some of these genes has already been described while that of many other genes among them IL7R and PCSK9 has not been reported yet.

**Table 2.** Breast cancer cell sensitivity to CD32-CR T cell-mediated cytotoxicity identifies overexpression of 42 genes associated with enhanced overall survival in advanced breast cancer patients.

| Gene name | log2FoldChange<br>expression of<br>sensitive vs resistant | Expression padj | Survival pvalue |
| --- | --- | --- | --- |
| MSN | 9,234237685 | 1,61E-28 | 0,0091 |
| ANXA1 | 5,680103532 | 3,59E-07 | 0,00012 |
| HLA-DRA | 5,553333035 | 1,54E-06 | 0,00023 |
| IL7R | 7,847016223 | 2,66E-06 | 0,017 |
| LIFR | 4,314342941 | 7,18E-06 | 0,0073 |
| PCSK9 | 7,797806132 | 9,48E-06 | 0,00046 |
| ICAM1 | 4,71910841 | 9,80E-06 | 0,0061 |
| SLC4A4 | 7,233043078 | 1,02E-05 | 0,031 |
| CD44 | 5,135518877 | 1,05E-05 | 0,000000034 |
| MBP | 5,863942435 | 1,05E-05 | 0,0091 |
| PDCD1LG2 | 5,564103601 | 0,000134633 | 0,0008 |
| CD38 | 3,655160914 | 0,000253909 | 0,0094 |
| ANTXR1 | 4,670855734 | 0,000265847 | 0,00014 |
| CUBN | 6,381850551 | 0,000307575 | 0,002 |
| PLAT | 6,696990062 | 0,000432311 | 0,00078 |
| FGFBP1 | 7,031398988 | 0,000463226 | 0,001 |
| HLA-DPB1 | 3,899732307 | 0,000538005 | 0,0028 |
| ITGB8 | 4,614246358 | 0,000589962 | 0,014 |
| LIPG | 5,903021878 | 0,001436942 | 0,00031 |
| HLA-A | 4,313972391 | 0,001450444 | 0,016 |
| HLA-DPA1 | 4,763693695 | 0,001638118 | 0,006 |
| SRPX2 | 4,126912066 | 0,002044177 | 0,00041 |
| NRROS | 5,079683458 | 0,005607296 | 0,032 |
| CDH2 | 3,75195309 | 0,005748377 | 0,018 |
| HBEGF | 4,941781375 | 0,00582612 | 0,012 |
| SLC6A2 | 8,393035966 | 0,006385352 | 0,00081 |
| BTN3A3 | 3,491327553 | 0,00694209 | 0,002 |
| CEACAM1 | 5,103805021 | 0,007057192 | 0,000002 |
| ITGA3 | 3,329192706 | 0,007409475 | 0,0069 |
| ITGB2 | 2,909231769 | 0,0095556 | 0,011 |
| SERPINE2 | 3,589186779 | 0,010257949 | 0,024 |
| ADTRP | 3,829816509 | 0,01390029 | 0,0038 |
| FAP | 5,764241237 | 0,021549094 | 0,00033 |
| DCBLD2 | 3,035651524 | 0,024155054 | 0,02 |
| PTPRC | 3,018702502 | 0,024542382 | 0,013 |
| CD22 | 3,04395917 | 0,024835513 | 0,00088 |
| HLA-H | 4,521019966 | 0,026629949 | 0,0045 |
| TNFRSF9 | 4,538715226 | 0,029923057 | 0,011 |
| MET | 3,95185859 | 0,030399118 | 0,0014 |
| IL15 | 3,471780313 | 0,036004979 | 0,024 |
| CD274 | 4,168703408 | 0,037513963 | 0,0037 |
| TRPV2 | 4,070284638 | 0,038141506 | 0,012 |

### Role of the ICAM1 in the regulation of the susceptibility of BC cells to the CD32A-CR T cell-mediated cytotoxicity

Among the forty-two overexpressed genes identified, we were particularly interested in the role of ICAM1 since it is involved in a variety of immunological functions including T cell binding, immunological synapse formation, and activation ^39^.

To validate the RNA-Seq study results, we assessed the distribution of ICAM1 protein expression among CD32A-CR sensitive and resistant cell lines by immunostaining with a PE-conjugated anti-CD54 mAb. Figure 4A shows that ICAM1 was preferentially expressed MDA-MB-468, MDA-MB-231, and HCC-1954 while resistant cells are negative or express it to a lesser extent (MCF-7, SK-BR3, and T-47-D) than sensitive cells. The well-defined ICAM1 binding to LFA1 might suggest that the disruption of this interaction could protect tumor cells from the cytotoxic effects of CD32A-CR T cells. Figure 4B shows that indeed, the addition to the cultures of an anti-CD18α mAb significantly reduced the susceptibility of BC cells to CD32A-CR T cells in a dose-dependent manner (dose range 1.25 μg/ml-10μg/ml). These results underline the key role of ICAM1/LFA-1 interaction in the generation of CD32A-CR T cell-mediated anti-tumor activity.

**Figure 4.**
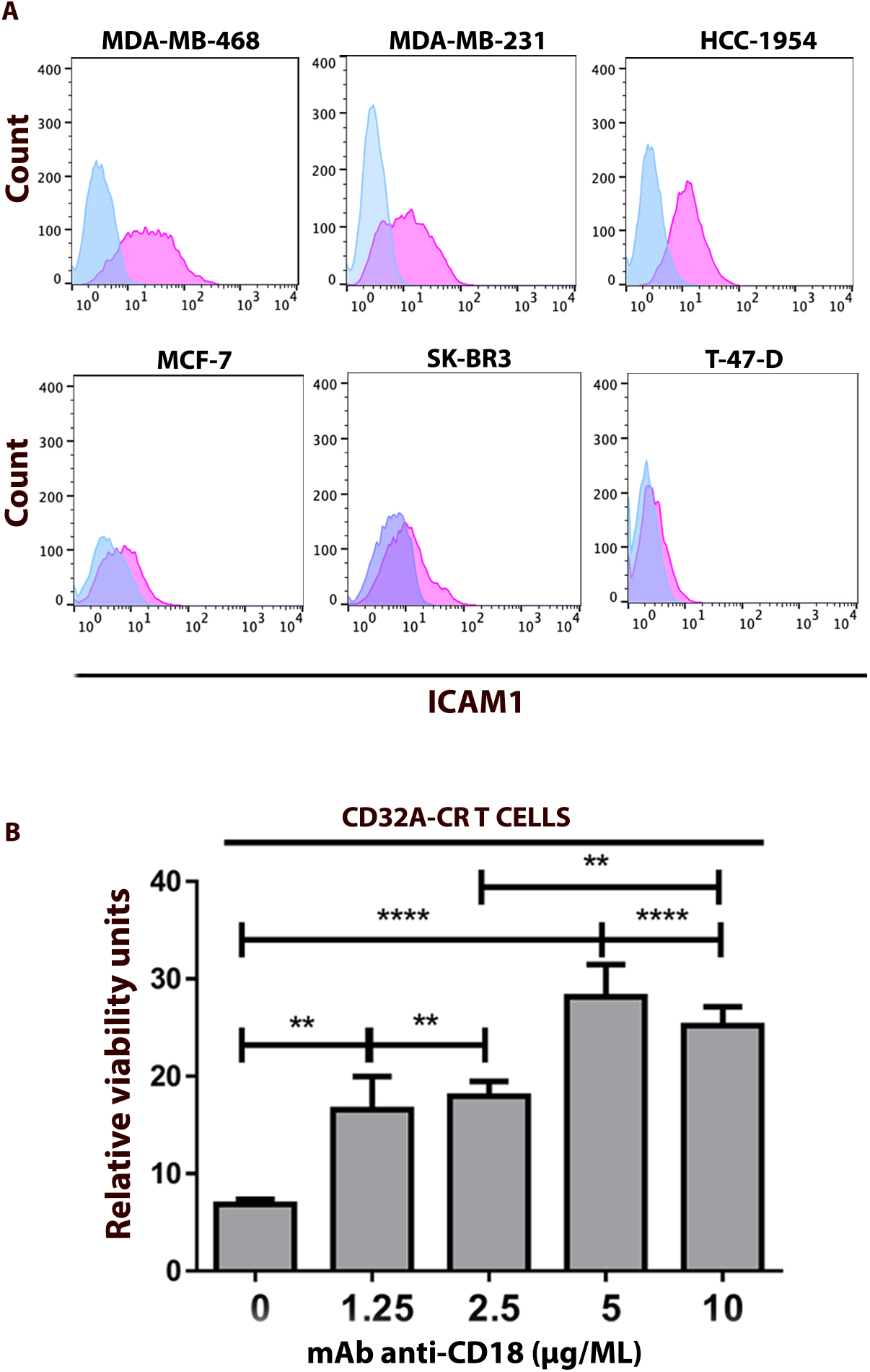
ICAM1 is preferentially expressed on the surface of sensitive BC cells and regulates CD32A-CR T cell anti-cancer activity. **A:** expression of ICAM1 on the surface of sensitive (upper panel) and resistant (lower panel) was assessed by immunostaining utilizing a Pe-conjugated anti-human ICAM1 (CD54). Following 30min incubation, cells were washed and analyzed by flow cytometry. **B**: rescue of MDA-MB-468 cell viability by LFA1-ICAM1 blockade was tested as follows: CD32A-CR T cells were pre-incubated with scalar concentrations of anti-CD18 mAb for 30min ant 37°C and washed. Then FcR negative MDA-MB-468 were added at an E:T ratio of 1.1. After 2 days, non-adherent cells were removed and the viability of MDA-MB-468 was measured by the MTT assay.

### Direct CD32A-CR T cell administration protects CB17-SCID mice from the MDA-MB-468 cell challenge

To address the *in vivo* anti-tumor potential of CD32A-CR T cells, three groups of CB17-SCID mice were injected subcutaneously with 1×10^6^ MDA-MB-468 cells alone, with panitumumab (150 μg) or together with 0.6×10^6^ CD32A-CR T cells, and tumor growth was monitored over time (fig. 5). CD32A-CR T cells, powerfully inhibited the TNBC breast cancer cell growth in 4 out of 4 mice (P = 0.01) while panitumumab had only a limited effect. As expected, while all untreated mice developed a measurable tumor within 30 days and had to be sacrificed within 100 days, only 1 out of 4 mice treated with panitumumab was disease-free and alive on day 120 of follow-up. In contrast, 4 out of 4 mice treated with CD32A-CR T cells were tumor-free and alive on day 120. These data indicate that CD32A-CR T cell-based immunotherapy is associated with a favorable course of the disease in the absence of therapeutic mAb administration.

**Figure 5.**
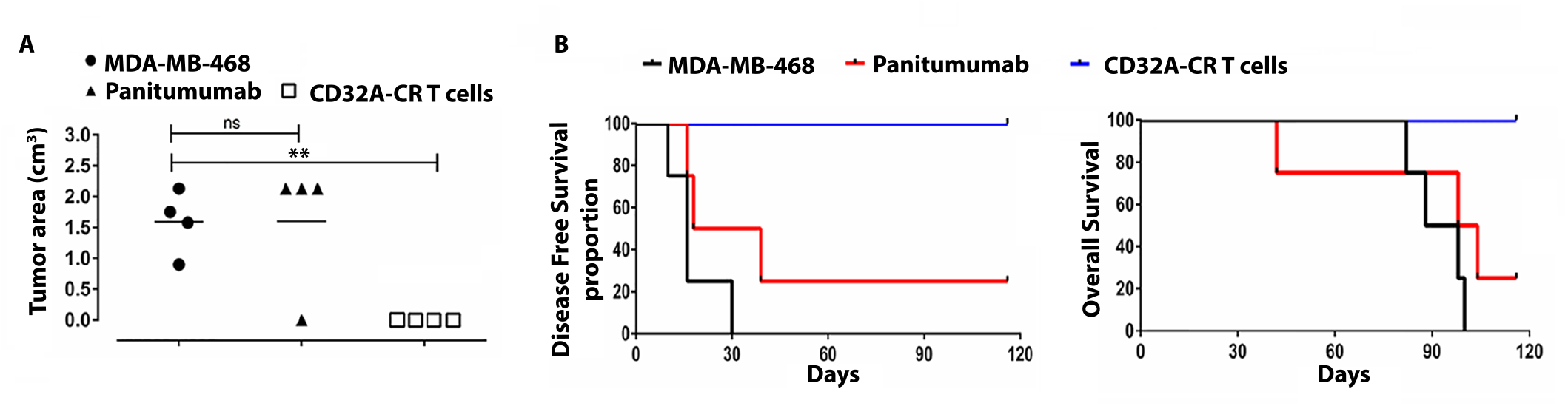
CD32A-CR T cells prevent tumor growth of CB17-SCID mice engrafted with MDA-MB-468 cells. Three groups of CB17-SCID mice (N =4 per group) received 1.0 ×10^6^ MDA-MB-468 cells subcutaneously, in the right flank, with or without panitumumab (150μg) or 0.6×10^6^ CD32A-CR T cells as shown. The left panel shows a scatterplot analysis of tumor volumes on day 120 post-injection. The middle and right panels show Disease-Free Survival and Overall Survival analysis. Data were analyzed by unpaired T-test **P < 0.01.

## DISCUSSION

CD16A, CD32A, and CD64 bind, with different affinities, to the Fc fragment of the Igs and pentraxins. The latter are a family of evolutionary conserved, soluble proteins present in the serum and mediate pro-inflammatory functions^13^. Interestingly, however, other than Fc and pentraxins, CD16 has been shown to bind putative surface ligand(s) on a variety of tumor cells leading to the activation of NK cells and cancer cell elimination^35^. However, likewise CD16, extracellular CD32A is composed of 2 immunoglobulin-like domains and binds with low affinity the Fc fragment of Igs and pentraxins ^12,34,40^. In the current state of knowledge, it is not known whether additional FcγRs recognize cell surface ligand(s) on pathological and/or healthy cells in the absence of target-specific antibodies.

Here, we have used CD32A-CR T cells to explore the localization of CD32A putative ligand(s) on the surface of 20 distinct cell types of which 15 were cancer cell lines. The molecular structure of CD32A-CR makes it a cytotoxic triggering molecule when it is transduced in cytotoxic cells. Therefore, crosslinking of CD32A-CR with its putative ligand(s), on the surface of the target cells, induces the activation of the T cell killing machinery. Concordantly, we have demonstrated that CD32A-CR T cells recognize CD32A-CR putative ligand(s), on MDA-MB-468 and MDA-MB-231 BC cells, resulting in specific CD32A downregulation and CD107A release consistent with cytotoxic granule exocytosis of T cells in an HLA-independent manner^41^. The direct anti-tumor activity is not restricted to the triple-negative BC cell lines but also involves HER2+ HCC-1954 cells, CRC HT29, and glioblastoma U-373MG, (data not shown) while HUVEC cells induced CD32A-CR downregulation. In contrast, no significant evidence of CD32A-downregulation is detectable upon conjugation with hematopoietic cells, a variety of epithelial cancer cells, two normal, primary fibroblasts, and one myoblast cell line, indicating that CD32A ligand(s) is preferentially expressed on BC cells.

The anti-tumor properties of CD32A-CR T cells have been confirmed by more than four sets of *in vitro* assays including confocal microscopy, bioluminescence, flow cytometry, and MTT assays. The observation that 3 out of 7 BC cell lines were sensitive to the engineered T cells suggests the existence of a subgroup of BC cells resistant to CD32A-CR T cell-mediated cytotoxicity.

The use of biosensors for the identification and localization of surface molecules with unknown ligand(s) or function is a reliable methodology and is consistent with our previous work in which bispecific monoclonal antibodies (BsAbs) allowed the identification of CD44 as a cytotoxic triggering molecule on NK cell and its metabolic pathways ^5,42,43^

One important issue is whether the direct cytotoxic activity of engineered T cells has any relationship with ADCC. We are comfortably convinced that CD32A-CR mediated cytotoxicity is an independent function since either human serum or purified IgGs failed to protect sensitive EGFR positive BC cells from CD32A-CR T cells, but rather their killing was enhanced in the presence of cetuximab. The results shown in the present study indicate that CD32A putative ligand(s) and Igs detain distinct docking sites on extracellular CD32A and support the role of CD32A-CR mediated cytotoxicity independent from ADCC. Also, BC cell killing has been achieved at a very low level of E:T ratio such as 0.5:1 even when the CD32A-CR T cell transduction has been lower than 100% indicating that the extent of CD32A-CR T cell anti-tumor activity is strikingly powerful.

Notably, we have observed that specific conjugation of CD32A-CR T cells with sensitive BC cells led to mutual cell killing while NT T cells did not. These results are consistent with published results indicating epithelial cancer cells and leukemia cells induced NK cell killing and CD16A downregulation^20,22,36^. Although the mechanism underlying this phenomenon is not readily apparent, it might be possible to hypothesize a feedback mechanism aimed to turn off an overwhelming immune response mediated by cytotoxic cells.

Although the anti-cancer activity of CD32A-CR T cells is preferentially directed towards BC cells, we have also evidence that some CRC and glioblastoma cell lines can be targeted (data not shown) while most hematopoietic malignant cells, we have utilized, are resistant. ^44^. Based on the susceptibility of cancer cells to CD32A-CR T cells, we have identified two populations of BC cells defined as sensitive or resistant. The RNA-seq analysis of the BC cell lines has underlined the compelling difference in gene expression between sensitive and resistant BC representing a remarkable transcriptomic signature for the identification of target cells. Among 42 differentially expressed genes, ICAM1 has come to our attention. ICAM1 is a key molecule involved in the leukocyte transmigration through the endothelial barrier but is also noteworthy that ICAM1 is heterogeneously expressed in BC tissues^45^. Therefore, the reduced susceptibility of sensitive BC cells to CD32A-CR T cell activity by the anti-CD18 mAb is likely to be due at least in part to the blockade of the interaction of β2 integrin on T cells^39^ with ICAM-1 leading to the inhibition of CD32A-CR T cell cytotoxicity.

Although ICAM1 appears to be required for the elicitation of an efficient CD32A-CR T cell anti-cancer activity (Figure 6) its expression is non-sufficient for the following considerations. First, optimal CD32A-CR direct cytotoxicity requires the presence of significant expression of ICAM1 and CD32A-CR ligand(s) as indicated by CD32A-CR profound downregulation in sensitive BC cells. Second, ICAM1 positive U937 and ML-2 cells have proven to be resistant to CD32A-CR T cells suggesting they lack CD32A-ligand(s). Third, the absence of ICAM 1 on resistant BC cells does not exclude the presence of CD32A-CR ligand(s) on these cells. In addition, the finding that ICAM1 positive HUVEC cells induced CD32A-CR downregulation is evidence of the presence of CD32A-ligand(s) on endothelial cells. In contrast, no significant CD32A-CR downregulation has been observed following cell-to-cell contact of CD32A-CR T cells with ICAM1 negative cells including HCT116 and A549.

**Figure 6.**
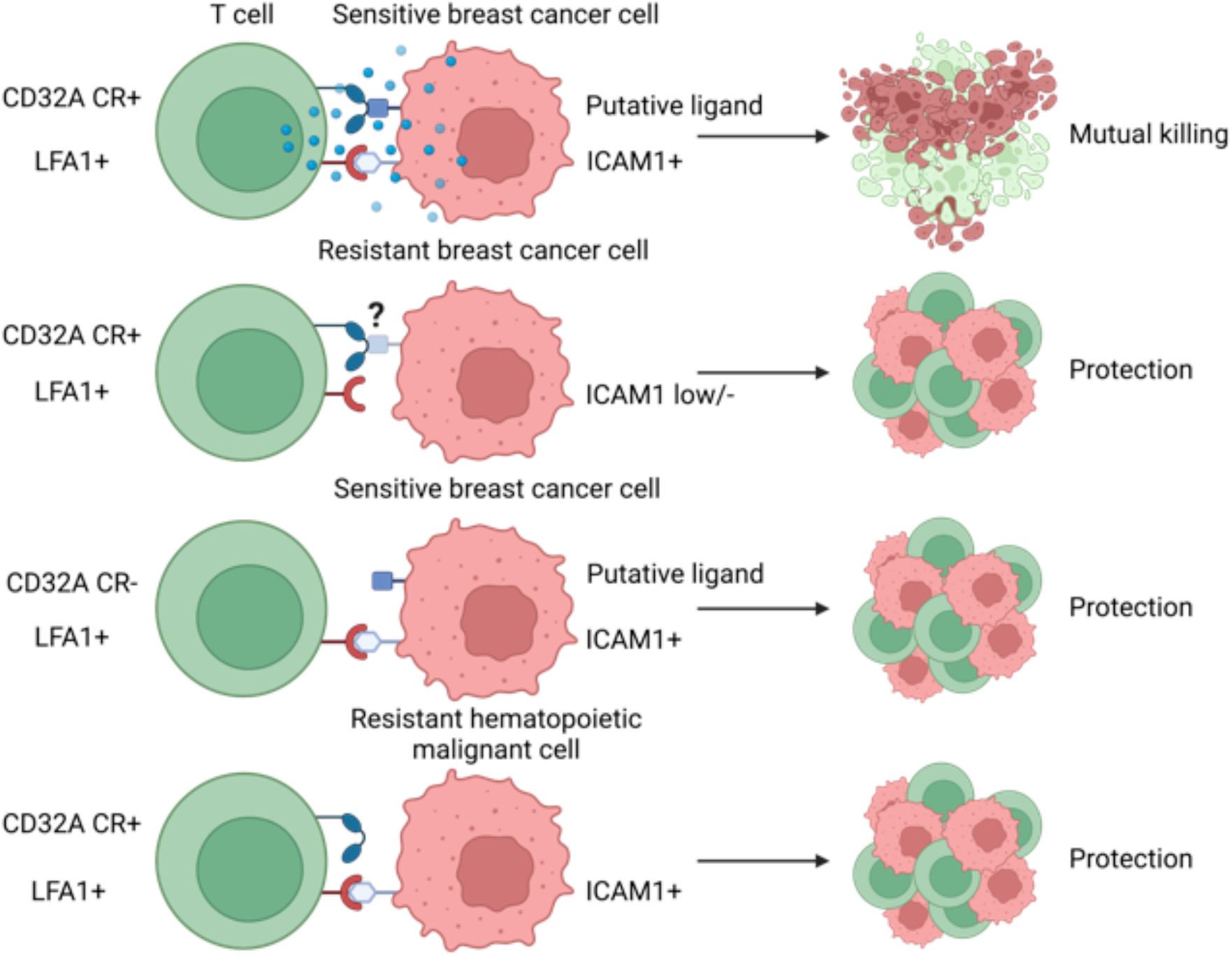
Hypothetical representation of cellular events occurring following conjugation between CD32A-CR T cells with sensitive or resistant cancer cells. **A:** optimal activation of CD32A-CR requires the expression of CD32A putative ligand(s) and ICAM1 on sensitive BC cells. Resistant BC cells are not susceptible to CD32A-CR T cells because they may express the ligand(s) but lack ICAM1 or both the ligand(s) and ICAM1. CD32A-CR negative NT T cells do not kill CD32A-CR sensitive BC cells, despite their expression of ICAM1 and putative CD32A ligand(s). Also, CD32A-CR positive T cells do not kill ICAM1+ hematopoietic malignant cells because of defective expression of CD32A CR ligand(s).

The information provided by this study raises some basic questions. What is the identity of CD32-CR ligand (s)? Is there any cross-talk between normal epithelial/endothelial cells and CD32A positive innate cells and what type of cellular response(s) are (is) generated? We are motivated to keep working on these issues since the availability of this information may have a relevant impact in the field of the biology of innate cells.

Nevertheless, despite the incomplete comprehension of the nature and distribution of CD32A putative ligand(s), CD32A-CR T cell-based immunotherapy, alone or in association with anti-EGFR therapeutic mAbs^32^, might be utilized for designing rational immunotherapies for targeting BC as suggested *in vivo* results.

Safety concerns are raised by the fact that HUVEC cells have induced significant downregulation of CD32A-CR. One may predict that engineered T cell immunotherapy may lead to endothelial cell damage reminding the capillary leak syndrome described during LAK cell-based and cluster differentiation mAb immunotherapy^46^ and in 19-CAR T cell immunotherapy^47^. However, similarly to classic CAR T cells, the safety of CD32A-CR T cells might be enhanced utilizing suicide genes. Unfortunately, an optimal animal model capable of providing reliable information about the safety of this type of immunotherapies is not available yet. It would be useful to determine whether the behavior of mouse and human FcγCR coincide allowing a more detailed investigation of CD32A-CR T cell off-shelf toxicity in syngeneic tumor-bearing mice models.

In addition, the identification of a gene signature associated with BC cell sensitivity to CD32A-CR T cells and predicting the favorable clinical course of BC in a retrospective study of 1400 patients with advanced disease opens new unsuspected investigation pathways in the field of innate cell and tumor biology.

## MATERIALS AND METHODS

### Antibodies and reagents

Allophycocyanin-conjugated mouse anti-CD3 (cat.555335), fluorescein isothiocyanate (FITC)-conjugated mouse anti-human CD3 (cat. 555332), FITC-conjugated mouse anti-human CD107a (cat. 555800), phycoerythrin (PE)-conjugated mouse anti-human CD32 (cat.550586), and CD54 (cat. 555511), purified NA/LE mouse anti-human CD3 (cat. 555329), purified NA/LE mouse anti-human CD28 (cat. 555725), purified mouse anti-human CD32 (8.26) (cat. 557333), purified mouse anti-human CD18 (cat. 556084) were purchased from BD Bioscience (San Jose, CA). Rabbit anti-human EGFR (D3B1) was purchased from Cell Signaling Technology (Danvers, MA, 01923). Cy-5-conjugated donkey anti-rabbit IgG and Alexa fluor-488 F(ab’)2 fragment goat anti-mouse IgG were purchased from Jackson ImmunoResearch Laboratories (Cambridgeshire, United Kingdom) and Invitrogen (Cambridge, MA, USA) respectively. Rabbit anti-asialo-GM1 (Cat-986-10001) was purchased from Wako Chemical Europe (Neuss, Germany)

Monensin (M5273), 3-(4,5-Dimethylthiazol-2-yl)-2,5-diphenyltetrazolium bromide (MTT), Hoechst nuclei staining, fluoromount aqueous mounting medium, and Dimethyl Sulfoxide (DMSO) were purchased from Sigma-Aldrich (Saint Louis, MO) and GeneJuice^®^ Transfection Reagent (Novagen) from Millipore (Burlington, MA). Human recombinant interleukin-7 (IL-7) and interleukin-15 (IL-15) were purchased from Miltenyi Biotec (Bergisch Gladbach, Germany). Retronectin (Recombinant Human Fibronectin) was purchased from Takara Bio (Saint-Germain-en-Laye, France). D-luciferin was purchased from PerkinElmer (Waltham, Massachusetts, USA). Dulbecco’s modified Eagle’s medium (DMEM), Iscove’s Modified Dulbecco’s Medium (IMDM), and Roswell Park Memorial Institute (RPMI) 1640 medium were purchased from Lonza (Basel, Switzerland). Fetal bovine serum (FBS), human AB serum, Phosphate-buffered solution (PBS), L-glutamine, and penicillin/streptomycin were purchased from Euroclone (Milan, Italy). Complete media (CM) were supplemented with 10% FBS, 2 mM L-glutamine, 0.1 mg/ml streptomycin, and 100 U/ml penicillin.

### Cell Lines

The packaging cell line 293T was cultured in IMDM CM and was used to generate helper-free retroviruses for T-cell transduction. Colorectal cancer (CRC) cell line HCT116 and HT-29 were maintained in RPMI CM. Non-small-cell-lung cancer (NSCLC) cell line A549, triple-negative breast cancer (TNBC) cells MDA-MB-231 and MDA-MB-468, SUM159 luminal A BC cells MCF-7 and T-47-D (ER^+^, PR^+/−^, HER2^−^), and HER2-enriched SKBR-3 and HCC-1954 (ER^−^, PR^−^, HER2^+^) were kindly provided by Dr. Antonio Rossi and Dr. Maria Lucibello (Institute of Translational Pharmacology, CNR, Rome, Italy). Submaxillary salivary gland A-253, Pharinynx squamous cell carcinoma FaDu, hematopoietic ML-2, U937, Jurkat, and human umbilical vein endothelial cell (HUVEC) were obtained from our laboratory cell collection. Human fibroblasts IMR-90 and BJ, and human myoblasts were kindly provided by Dr. Lucia Latella (Institute of Translational Pharmacology, CNR, Rome, Italy) Table 1. All the cell lines were mycoplasma-free and were maintained in DMEM CM. Human myoblasts were grown in DMEM supplemented with Primary Skeletal Muscle Growth Kit (ATCC). MDA-MB-468-Luc+ cells were transduced at a multiplicity of infection (MOI) 10 with a third-generation self-inactivating lentiviral vector expressing firefly luciferase as previously described^48^. Quantification of light emission generated by luciferase-expressing cells through ATP-dependent conversion of luciferin to oxyluciferin can be used as a nonlytic, real-time cell viability assay.

### CD32A-chimeric receptor production and T-cell transduction

The construction of the CD32A^−^chimeric receptor (CR) [patent: Priority Number n. 102014902258369 (exTO2014A000361_06/05/2014)], virus production, and T cell transduction have previously been described^32^. Briefly, HEK 293T cells were cotransfected with CD32A-CR-SFG, PegPam, and RDF vectors. After 48 and 72hr, cell supernatants containing viral particles were harvested and stored at −80°C. Human peripheral blood mononuclear cells (PBMCs) were isolated from buffy coats and seeded in 24-well plates precoated with anti-CD3 and anti-CD28 mAbs for 48hr. Activated T cells were collected and incubated with retroviral particles carrying CD32A-CR for 72hr into a retronectin-coated 24 well-plate. After transduction, T cells were kept in culture for 12–15 days before using for further experiments. Transduced and Non-Transduced T cells were maintained in RPMI CM supplemented with 5ng/ml IL15 and 10 ng/ml IL7

### Confocal Microscopy

CD32A^−^CR–T cells were cocultured with MDA-MB-468 breast cancer cells at an effector: target (E:T) ratio of 2:1 on poly-d-lysine (0.02%) pre-coated glass multi-chambers well. After a 2hr incubation, cells were fixed with a solution containing 4% of paraformaldehyde (PFA). Samples were then washed and incubated with an Fc gamma blocking reagent to avoid nonspecific staining. Cells were incubated with rabbit anti-human EGFR and mouse anti-human CD32 antibodies (1:50 dilution) for 1h at room temperature in a humidity chamber. Cells were then stained with cy-5-conjugated donkey anti-rabbit and alexafluor-488 goat anti-mouse secondary antibodies, while cell nuclei were counterstained with Hoechst for 5 min at RT. Samples were acquired using a 63x oil immersion lens on a Leica Laser Scanning confocal microscope (Leica Microsystems).

### In Vitro Tumor Cell Viability Assay

Antitumor activity of CD32A-CR T cells was evaluated in vitro by a cell viability assay using an MTT reagent. Tumor target cells (20 × 10^3^/well) were seeded in triplicate in 96-well plates, and CD32A-CR or non-transduced T cells were added at different E:T ratios. Following a 12-hr incubation at 37°C, non-adherent T cells were removed and the remaining adherent target cells were incubated with 100 μl of fresh medium supplemented with 20 μl of MTT (5 mg/ml) for 3hr at 37°C. Supernatants were then removed and 100 μl of DMSO were added to each well and placed on a plate shaker for 15 min protected from light. Absorbance was measured at 570 nm.

For in vitro Bioluminescent Imaging (BLI), MDA-MB-468 luciferase-expressing cells were seeded in 96-well microplates in triplicate in presence of CD32A-CR T cells or non-transduced cells as a negative control for 72hr at 37°C at different E: T ratios. Cell culture medium was then supplemented with D-luciferin (PerkinElmer) dissolved in PBS (150 μg/mL) 10min before analysis performed using the IVIS^®^ Lumina II platform (PerkinElmer, Waltham, MA, USA). Photons emitted from luciferase-expressing cells in selected regions of interest (ROI) were quantified using the Living Image^®^software.

### CD107a Assay

CD107a release was tested by incubating CD32A–CR T cells with A549, HCT 116, MDA-MB-468, and MDA-MB-231 target cells at an E:T of 2:1 for 1hr at 37°C in a 5% CO_2_. FITC mouse anti-human CD107a antibody diluted 1: 50 was added in a final volume of 200 μl in a 96 multi-well plate. Then, 2 μM of monensin (Golgi stop) was added to each well and cell coculture was incubated for 4hr at 37°C. After incubation, samples were stained with PE mouse anti-human CD32 for 30min at 4°C, washed, and fixed in 1% PFA. The CD107a expression on CD32A-CR T cell surface was assessed by using a 2-laser BD FACSCalibur flow cytometer (Becton Dickinson, Franklin Lakes, NJ). Results were analyzed using Tree Star, Inc. FlowJo software.

### RNA-Seq Analysis of Breast Cancer Cell Lines

RNA-Seq from 6 breast cancer cell lines was downloaded from the NCBI SRA portal as part of the Cancer Cell Line Encyclopedia (SRP186687) project and transformed into fastq files. All the preprocessing steps to trim adapters and remove low-quality reads were performed with the dedicated fastp ^49^ tool version 0.21.0. GRCh37/hg19 human reference genome was used for all analyses. Fastq files were aligned to the reference genome using HISAT2 ^50^ version 2.2.1 and parameters --no-mixed --no-discordant --very-sensitive --known-splicesite-infile --dta-cufflinks -x genome_snp_tran. Alignments in SAM format were converted in the binary BAM format, sorted by genomic coordinates, and indexed by SAMtools. Gene expression quantification was obtained with the featureCounts tool of the Subread ^51^ 2.0.2 package (parameters ‘--countReadPairs’ and ‘-p’) and the gencode annotation v38lift37. Counts' normalization and differential gene expression analysis was performed using the well-known Bioconductor package in R DESeq2 ^**52**^ version 1.32.0 following the manufacturer’s instructions. Gene set enrichment analysis was performed using the ‘Investigate Gene Set’ tool from the GSEA web portal (*https://www.gsea-msigdb.org*) and the GO Cellular Component ontology. Plotting and statistics were done using specific R packages.

### Xenograft Mouse Model

In vivo experiments were performed following European Directive 2010/63/ EU guidelines and regulations. The Italian Ministry of Health approved animal handling and procedures (authorization code: 186/2016-PR). Antitumor activity of CD32A-CR T cells was assessed using 8-week-old male CB17-SCID mice (CB17/lcr-PrkdcSCID/lcrlcoCrl, Charles River Laboratories, Lecco, Italy, Cat. CRL:236, RRID: IMSR CRL:236), 12-18 g body weight, engrafted with MDA-MB-468 TNBC cells. Mice were housed in temperature-controlled rooms with a 12hr light/dark cycle and free access to sterile water and autoclaved standard chow diet (4RF25; Mucedola, Milan, Italy). Endogenous NK cell activity was suppressed by intraperitoneal injection of a 20 μl rabbit anti-asialo-GM1 antibody. Mice received anti-asialo-GM1 antibody on days −3, 0, +14, and +21 since tumor cell engraftment. On day 0, mice were grafted subcutaneously in the right flank, with 1 × 10^6^ MDA-MB-468 cells, and then randomly separated into two groups (4 mice per group). Group 1 received only tumor cells and group 2 the effector cells and tumor cells at an E:T ratio of 0.6:1. Every 3 days, tumor volumes (TV) were measured with caliper and calculated using the formula: TV (cm^3^) = 4/3pr^3^, where r = (length + width)/4. Mice were sacrificed when tumor volume reached 2 cm^3^.

## STATISTICAL ANALYSIS

The unpaired t-test or the Mann–Whitney test were used to analyze the results. Survival time differences were determined using the Kaplan-Meyer method and the log-rank-(Mantel-Cox) test. Differences with a p-value <0.05 were considered statistically significant.

## DATA AVAILABILITY

The data supporting the findings of this study are available from the corresponding author upon reasonable request.

## ACKNOWLEDGMENTS

GS is supported by the Italian Association for Cancer Research (AIRC) Foundation: Investigator Grants (IG) 2015-17120 and (IG) 2020-24440.

We thank Dr. Mauro Cozzolino and Aymone Gurtner for confocal microscopy technical assistance and Dr. Antonio Rossi for the helpful discussion. We also thank Dr. Pamela Papa and Dr. Matilde Paggiolu for administrative assistance.

